# Exploiting attP landing sites and gypsy retrovirus insulators to identify and study viral suppressors of RNA silencing

**DOI:** 10.1101/2025.02.12.637972

**Authors:** Adarsh K. Gupta, Pratima R. Chennuri, Raquel D. Monfardini, Kevin M. Myles

## Abstract

RNA interference (RNAi) pathways are crucial for regulating viral infections in both animals and plants, acting as defense mechanisms that limit pathogen replication. This has led to the evolution of viral suppressors of RNA silencing (VSRs) across various plant and insect viruses, with potential analogs in arthropod-borne human pathogens. However, while functionally similar, VSRs often lack genetic conservation due to convergent evolution. Research on VSRs typically involves analyzing individual proteins expressed in host cells with secondary reporter constructs, but the lack of a standardized system can lead to inconsistent findings. Our study examined how genomic insertion sites affect VSR activity using a transgenic *Drosophila melanogaster* reporter system. We integrated the VSR protein DCV-1A into three different attP sites and assessed silencing. The results showed significant variation in VSR activity across loci due to position effects. However, by flanking the transgenes with gypsy retrovirus insulators, we achieved consistent high-level silencing across all sites. These findings suggest the potential for establishing a standardized reporter system in *Drosophila*, facilitating the identification, study and comparison of VSR proteins. However, our results also highlight the limitations of using isolated proteins in reporter systems, emphasizing the need for a comprehensive holistic approach to definitively determine VSR functions.

## INTRODUCTION

RNA interference (RNAi) pathways play a pivotal role in regulating the virulence of viruses across both animal and plant kingdoms [1–4]. These pathways act as defense mechanisms, limiting pathogen replication and thereby prompting the evolution of countermeasures by pathogens [5,6]. Viral suppressors of RNA silencing (VSRs) have been detected in various plant virus families and several insect viruses [6,7]. Additionally, certain sequences and proteins present in arthropod-borne human pathogens are speculated to function as VSRs [5,8]. These ongoing evolutionary arms races have independently led to the emergence of similar VSR strategies on multiple occasions [5,9]. Consequently, while VSRs may share functional characteristics at the biochemical level, they often display minimal genetic conservation [9].

Research on potential VSRs typically focuses on the analysis of individual proteins, which are expressed either transiently or stably within host cells, alongside a secondary reporter construct to measure their activity [10–13]. The absence of a uniform reporter system, however, has led to the use of a broad array of cell types, promoters, and transgene reporters to evaluate VSR function. This diversity in experimental approaches can lead to false-positive or -negative results, as well as inconsistent and non-replicable findings across different research systems. Although stable genetic integration (transgenic expression) offers certain benefits over temporary transient expression, including the potential to establish a uniform testing protocol, the expression levels of VSRs may still be influenced by the genomic context of their insertion site in the host DNA, known as position effects [14,15].

The aim of our study was to examine how genomic insertion sites influence the expression of candidate VSR proteins [16–18] using a transgenic *Drosophila melanogaster* reporter system [11]. Our findings reveal that silencing varies depending on the phiC31 docking site [19–21], suggesting that insufficient expression from specific genomic locations within *Drosophila*, and potentially other organisms, may impede the detection of VSR activity. However, we demonstrate that integrating gypsy insulators [22,23], which are capable of significantly enhancing gene expression in comparison to the uninsulated site [24], can mitigate these issues. These results pave the way for establishing a standardized reporter system in *Drosophila* with consistent levels of transgene expression, facilitating the initial assessment of viral proteins and enabling meaningful comparisons of VSR activity. Moreover, this information may be used to develop better reporter systems in other organisms.

## RESULTS

### Tissue-specific induction of a VSR and position effects at attP docking sites

We generated transgenic flies that express the well-characterized VSR protein, *Drosophila* C virus (DCV)-1A [18], using the binary GAL4-UAS [25] system (fig. 1 and s1). Conditional expression in the *Drosophila* eye was achieved (fig. 1) with a GMR-GAL4 driver [26,27]. To systematically evaluate the effects of host chromatin position on VSR protein expression, a UAS-DCV-1A cassette (fig. 1 and s1) was integrated into three different attP landing sites: VK1 [21], attP40 [19], and attP18 [20].

**Figure 1.**
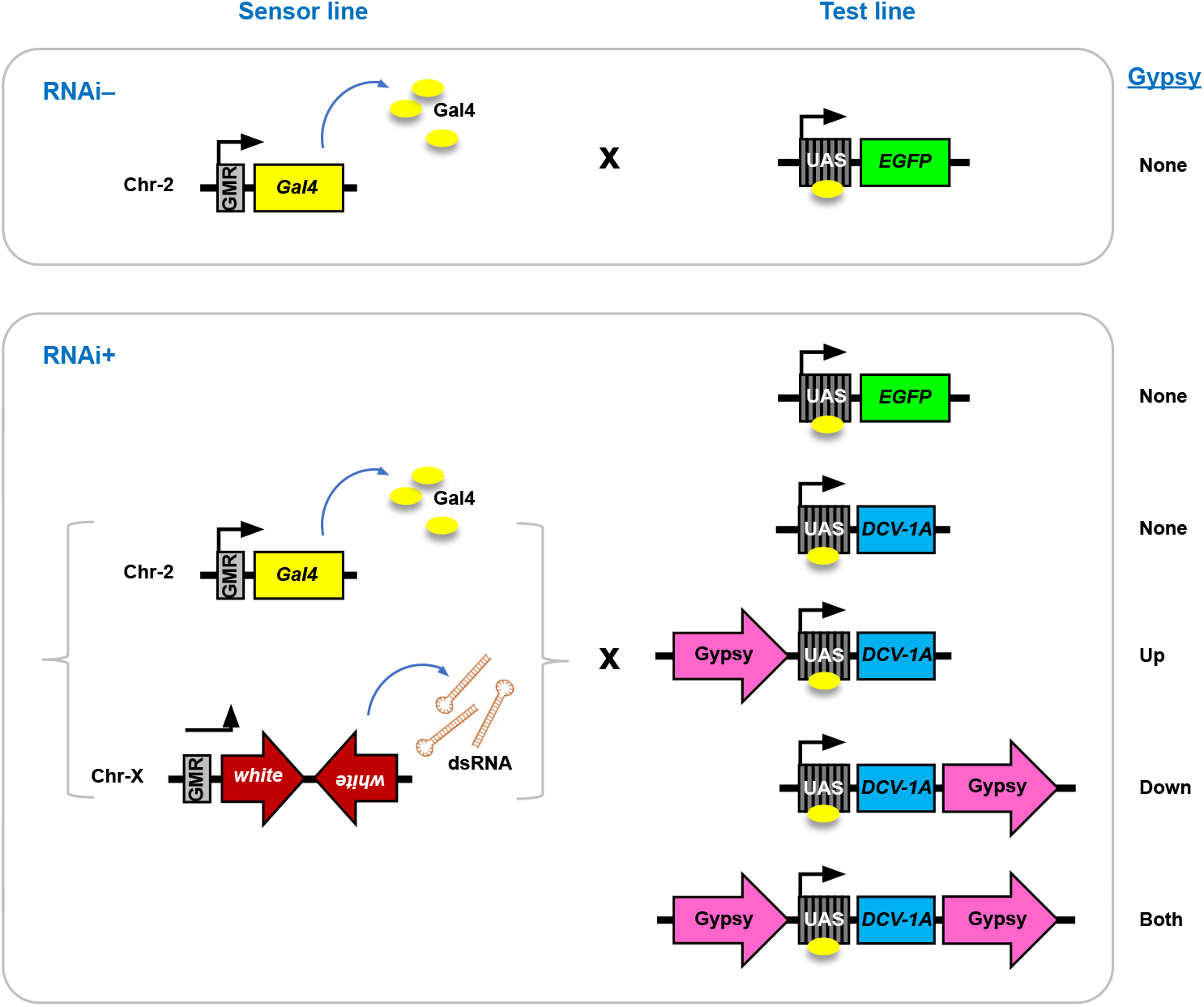
A transgenic reporter system designed to silence the *Drosophila white* gene using a dsRNA hairpin sequence. In the sensor lines, both the *white* inverted repeat sequence and the yeast GAL4 protein are regulated by the GMR enhancer, which is active in immature and adult retinal tissue where the endogenous white gene is also expressed. The tissue-specific expression of the inverted repeats produces hairpin-loop RNA, inducing RNAi that targets the *Drosophila white* gene. In the test lines, VSR proteins or controls (such as EGFP) are under the transcriptional control of a *Drosophila* promoter linked to GAL4-responsive upstream activating sequence (UAS) repeats. When the test lines are crossed with reporter lines, GAL4 drives the tissue-specific expression of VSR or control sequences. Additionally, some test lines were created with gypsy insulator sequences positioned either upstream, downstream, or flanking the GAL4-responsive elements.

In order to assess position effects at these attP landing sites using a biologically relevant assay, the VSR-expressing lines were crossed with an RNAi-sensor line (fig. 1). The RNAi-sensor contains two transgenic cassettes, GMR-*white* inverted repeat (IR) on the X chromosome and GMR-GAL4 on chromosome 2 (fig. 1). The *Drosophila white* gene encodes an ABC transporter crucial for eye pigmentation [28], allowing its expression to be phenotypically monitored and indirectly quantified in terms of eye pigment levels. The *white*+ eye phenotype is dark red (fig. 2), while a *white*- or null mutant displays white eyes. Silencing of the white target gene mediated through the expression of a double-stranded RNA hairpin from the GMR-Gal4-*white*-IR transgene [11] results in an intermediate or orange phenotype, with paleness depending on the level of silencing (fig. 2).

**Figure 2.**
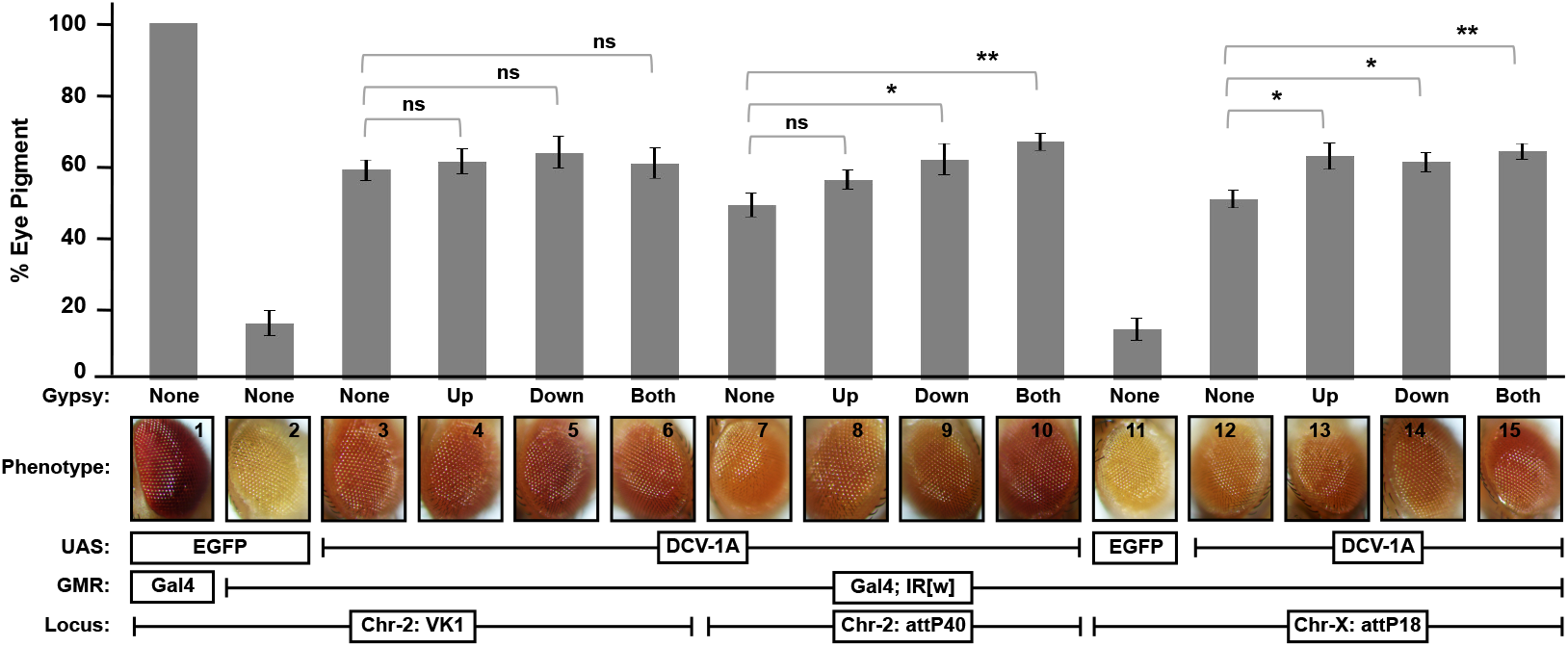
Position effects and DCV-1A mediated suppression of silencing by the *white* GMR-hairpin transgene in the presence or absence of gypsy insulators. Far left, (1) eye color phenotype of GMR-GAL4; UAS-GFP flies (VK1). (2) Eye color of flies bearing GMR-*white*IR; GMR-GAL4; UAS-GFP transgenes (VK1). (3-6) Suppression of *white* silencing by the GMR-hairpin by DCV-1A expressed from the VK1 landing site with upstream, downstream, flanking (both), or no gypsy insulators. (7-10) Suppression from the attP40 landing site with upstream, downstream, flanking, or no gypsy insulators. (11) Eye color phenotype of GMR-*white*IR; GMR-GAL4; UAS-GFP (attP18) flies. (12-15) Suppression from attP18 landing site with upstream, downstream, flanking, or no gypsy insulators. Eye pigment levels were measured in three separate experiments using age-matched 4-day-old adults, and are expressed as a percentage relative to the pigment levels measured in the GMR-GAL4; UAS-GFP (VK1) controls. Error bars represent standard error. Significance was calculated by Student’s *t*-test.

In order to determine the landing site permitting suppression of target gene silencing to the greatest degree, we quantified eye pigment levels in GMR-GAL4-*white*-IR adults expressing DCV-1A from the various attP loci (fig. 2). Notably, we found that expression from the VK1 locus on chromosome 2 suppressed silencing of the target gene by the greatest amount (fig. 2). Expression of DCV-1A from the other two landing-site loci-attP40 and attP18-resulted in less robust suppression than was observed at the VK1 locus (fig. 2). These results suggest that position effects may influence the results of phenotypic assays measuring VSR activity.

### Gypsy insulators facilitate optimal VSR expression at multiple attP landing sites

Although the above results showed that the VK1 attP site exhibits the highest level of VSR expression among the landing sites evaluated (fig. 2), position effects might still prevent optimal expression from this locus. Flanking transgenes with insulators has previously been shown to block the effects of neighboring enhancers, silencers, as well as heterochromatin (i.e., position effects), in order to ensure that expression of the transgenes is sufficient to produce a detectable phenotype [22]. The gypsy retrovirus insulator in particular has been previously found to boost gene expression to levels several fold greater than un-insulated loci [24]. Thus, in order to test if VSR expression could be further optimized, we created additional transgenic UAS-DCV-1A lines with flanking gypsy retrovirus insulators.

Flanking gypsy insulators did not affect VSR activity at the VK1 locus (fig. 2), suggesting that the expression of DCV-1A at this landing site cannot be optimized further. However, we found that flanking the UAS-DCV-1A construct with gypsy insulators equalized VSR activity across all three landing sites, significantly increasing suppression at attP40 and attP18 loci (fig. 2). These results suggest that gypsy insulators can be used to express VSR proteins at consistent and high levels across permissive loci.

### Transgenic RNAi sensor reporter systems sometimes fail to identify well characterized VSRs

Consistent levels of VSR expression from transgene constructs with flanking gypsy insulators (fig. 1) at specific genomic integration sites (fig. 2) suggest that it is possible to compare the activity of VSR proteins from different viruses (fig. 3). Therefore, we compared the VSR activity of DCV-1A at the VK1 attP site with that of another well-characterized VSR protein, flock house virus (FHV) B2 [16,29–32], expressed from a transgene (UAS-FHV B2) integrated at the same locus. Both transgenes contained flanking gypsy insulators to ensure optimal expression [24], as position effects may also depend on the inserted transgene sequence. Surprisingly, FHV-B2 did not exhibit any evidence of VSR activity (fig. 3). After confirming the sequences of both B2 and the regulatory elements present in the integrated UAS-FHV B2 cassette (fig. S1 and S2), we assessed VSR activity at the attP18 landing site. Consistent with the results reported above, no evidence of VSR activity was observed at the attP18 locus either (fig. 3). To confirm that our transgenic reporter system could identify VSR proteins other than DCV-1A, we generated transgenic flies expressing a third well-characterized VSR, Cricket Paralysis virus (CrPV) 1A [17]. Significant suppression of target gene silencing was observed in GMR-GAL4-white-IR adults expressing CrPV-1A (fig. 3). Finally, expression of the B2 protein from the UAS-FHV B2 transgene integrated at both the VK1 and attP18 landing sites was confirmed through western blotting (fig. S3). These results illustrate the limitations of studying VSRs as isolated proteins with reporter systems.

**Figure 3.**
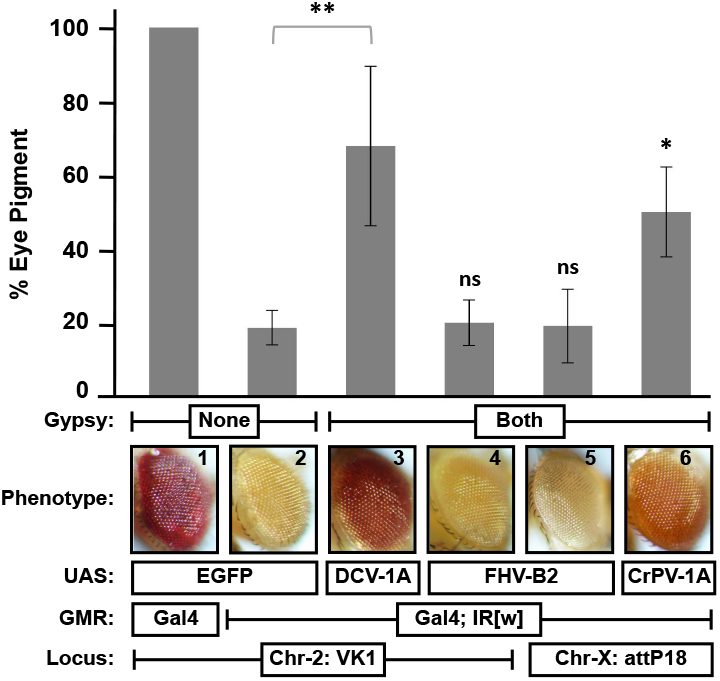
Suppression of GMR-hairpin mediated silencing of *white* by DCV-1A, CrPV-1A, or FHV-B2. Far left, (1) Eye pigmentation of GMR-GAL4; UAS-GFP flies (VK1). (2) Eye color observed in flies with GMR-*white*IR; GMR-GAL4; UAS-GFP transgenes (VK1). (3) Suppression of *white* silencing mediated by DCV-1A expressed from the VK1 landing site with flanking gypsy insulators. (4-5) Lack of *white* silencing inhibition by FHV-B2 expressed from the VK1 or attP18 landing sites with flanking gypsy insulators. (6) Suppression of *white* silencing by CrPV-1A expressed from the attP18 landing site with flanking gypsy insulators. Eye pigment levels were measured in three separate experiments using age-matched 4-day-old adults, and are expressed as a percentage relative to the pigment levels measured in the GMR-GAL4; UAS-GFP (VK1) controls. Error bars represent standard error. Significance was calculated by Student’s *t*-test.

## DISCUSSION

The attP landing sites used in this study were chosen because they have in previous work been associated with high levels of gene expression [19–21,33]. Thus, position effects may prevent optimal expression to an even greater degree at other loci or tissues, suggesting that each locus must be selected on a case-by-case basis after testing. Additionally, the proteins used in this study have been associated with robust VSR functions [16–18,29–32]. Lower-level expression of proteins with less robust VSR activity could further hinder positive identification with reporter assays. However, we show here that consistently high levels of suppression can be achieved at permissive loci (fig. 2) by flanking VSR expressing transgenes with gypsy retrovirus insulators [24]. An advantage of this system is that it ensures consistent levels of VSR expression across well characterized loci on multiple chromosomes (fig. 2). The ability to create lines with consistently high and equivalent levels of transgene expression is important for using reporter systems to identify, study, and compare VSR proteins.

However, our studies also highlighted the limitations of studying isolated proteins with reporter systems. The VSR function of B2 has been conclusively demonstrated in genetic rescue experiments, considered the gold standard for identifying such proteins. The replication of a B2-defective FHV has been rescued in both dcr-2 and ago-2 null mutants, unequivocally implicating the protein as a VSR [16,32]. However, examining the protein in isolation with a trans-acting reporter assay did not identify the VSR function previously associated with B2 (fig. 3). While some studies have previously shown the VSR activity of B2 in reporter assays [12,34], our inability to detect this function highlights the inherent difficulties of comparing results from different reporter assays. Our study is also not the first to show that reporter assays sometimes fail to detect the VSR activity of proteins with VSR functions [9,34].

Nevertheless, viral proteins are often multifunctional, and it is not always possible to obtain viable mutants knocking out only VSR functions, precluding genetic rescue experiments [35,36]. Therefore, reporter assays will continue to be important tools for identifying and studying VSR proteins. However, our findings underscore the importance of taking a comprehensive, holistic approach to definitively identify or rule out authentic VSR functions. The results of reporter assays should not be considered in isolation as definitive determinations of VSR function or lack thereof.

## MATERIALS AND METHODS

### Transgenic flies and genetic crosses

A balanced transgenic sensor line of RNAi (P{GMR-w.IR}; P{w[+mC] GMR-GAL4}/CyO) was generated through genetic cross of two transgenic lines: Bloomington Drosophila Stock # 32068 (P{GMR-w.IR}13D; P{ry[+t7.2]=neoFRT}40A r2d2[S165fsX] [11] and 1104 (w[*]; P{w[+mC]=GAL4-ninaE.GMR}12). All transgenic test lines were engineered with transgenes (EGFP, DCV-1A, FHV-B2 or CrPV-1A) to express under Gal4-inducible UAS promoter [25], with (up, down or both) or without gypsy elements (fig. 1). UAS constructs (fig. S1) were integrated into *D. melanogaster* genome via ϕC31-integrase mediated recombination [33,37] at VK1 [21] or attP40 landing site [19] on Chr-2, or at attP18 landing site [20] on Chr-X (GenetiVision Corporation, Houston, TX). Experimental crosses (fig. 1) were performed between age-matched sensor (female) and test (male) flies, followed by progeny collection at 4 days-post-eclosion for further analyses.

### Eye pigment assays

Eye pigment extractions were performed as described [34] with few modifications. Heads of 5 virgin female adults at 4 days-post-eclosion were homogenized in acidified methanol (0.1% HCl), followed by gentle rocking for 24 hrs at 4°C. After pigment extraction, samples were incubated at 50°C for 5 min, followed by high-speed centrifugation. Optical density of each clarified supernatant sample was read as absorbance at 480 nm.

### Western blotting

Total proteins were extracted by homogenizing 5 heads of virgin female flies at 4 days-post-eclosion in ice-cold protein extraction buffer [250 mM HEPES-KOH, 100 mM KCl, 10 mM EDTA, 0.1% Triton X, 5% Glycerol with freshly added 5 mM DTT and Protease Inhibitor Cocktail (Roche, Indianapolis, IN, USA)]. The supernatant was resolved by SDS-PAGE analysis, followed by dry-transfer onto a PVDF membrane as described [38] using iBlot 3 System (Invitrogen). Protein blot was incubated in 5% (w/v) Blotting-Grade Blocker (Bio-Rad, Hercules, CA, USA), followed by FHV-B2-specific polyclonal monospecific rabbit antiserum (Pacific Immunology, San Diego, CA) or β-actin-specific polyclonal rabbit antibodies (Thermo Scientific, Waltham, MA, USA) at 1:10,000. Goat anti-rabbit HRP conjugate (Thermo Scientific) was used as a secondary antibody at 1:25,000. Immunoblots were developed using Immobilin Western Chemiluminescent HRP Substrate (Millipore, Billerica, MA, USA), followed by visualization of signals with iBright CL1500 Imaging System (Thermo Scientific).

## ACKNOWLEDGEMENTS

We thank Snigdha Musugunthan for her assistance in rearing and maintaining *Drosophila melanogaster* stock lines used in this study.

## AUTHOR CONTRIBUTIONS

KM, AG, and PC conceptualized the study; KM, AG, and PC designed the research; AG, PC, RM and KM performed the experiments and analyzed the data; KM and AG wrote the manuscript; KM, AG, and PC revised the manuscript. All authors approved the final manuscript.

## FIGURE LEGENDS

**Figure S1.**
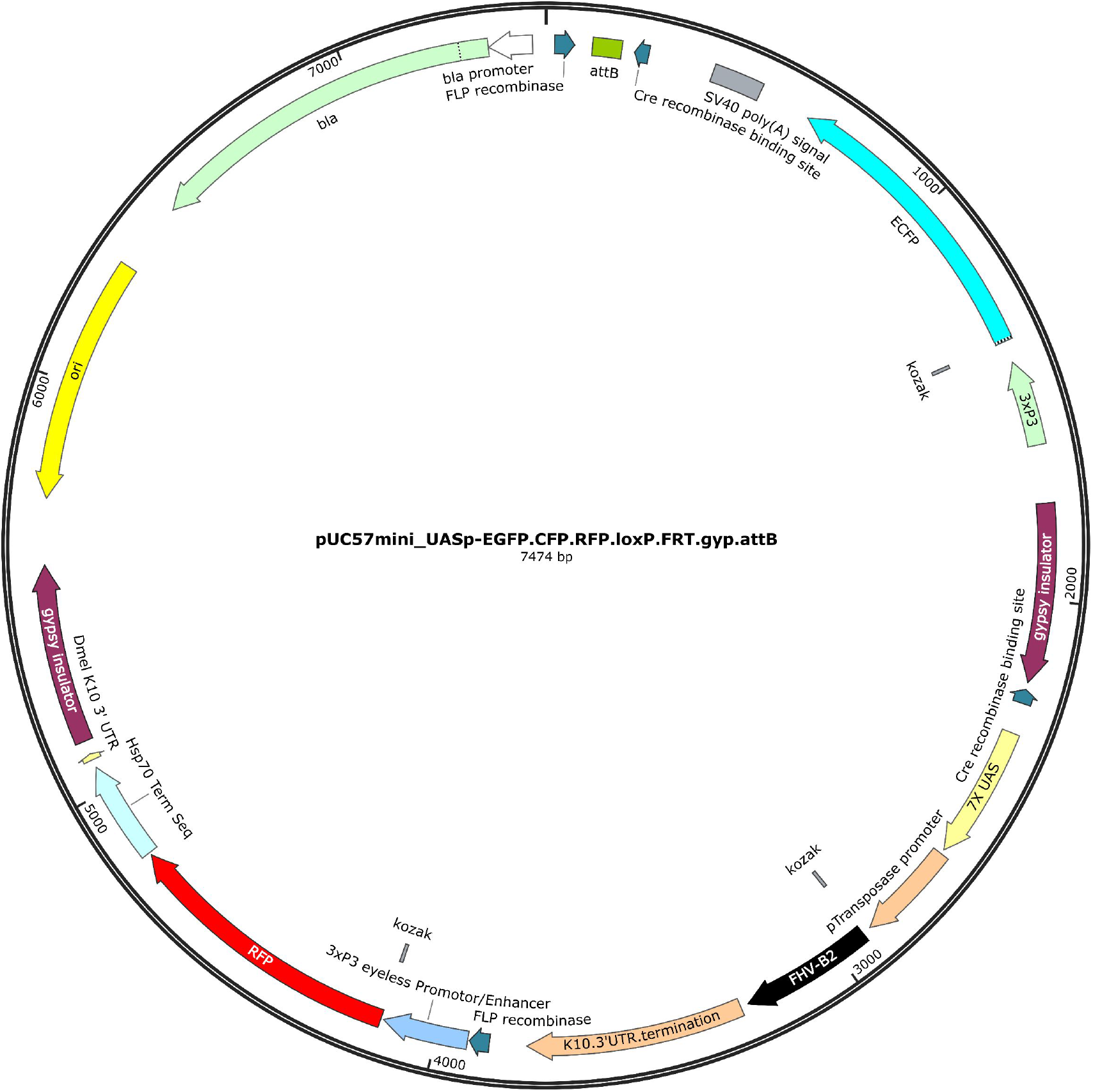
Map of UAS expression vector. Vector with FHV-B2 (black) and flanking gypsy elements (maroon) is shown here as an example. The full sequence of this construct has been deposited in the GenBank database under the accession number OR769027.

**Figure S2.**
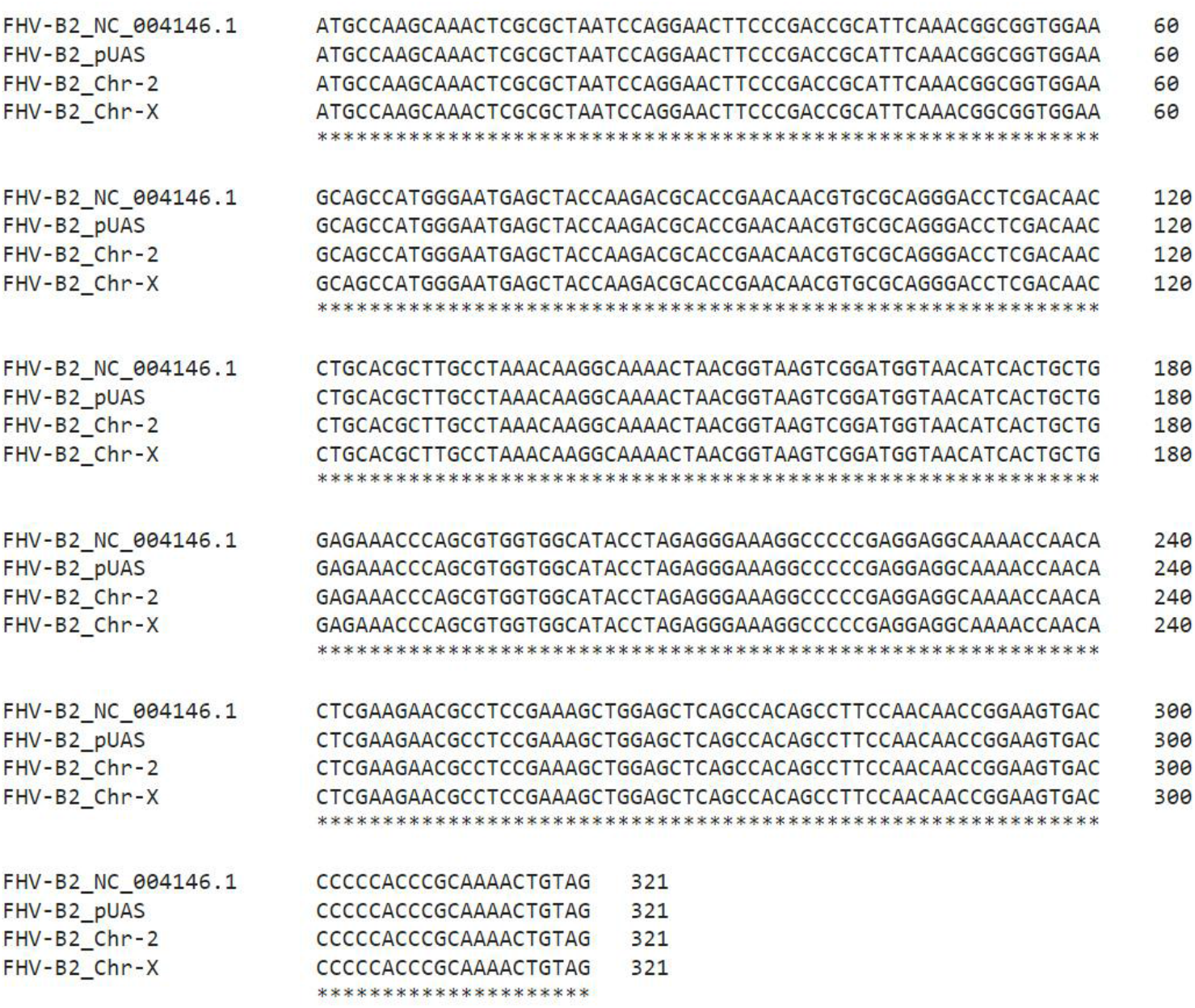
Confirmation of sequence accuracy for UAS-FHV-B2 vector. The FHV-B2 reference sequence was obtained from GenBank, accession no. NC_004146.1.

**Figure S3.**
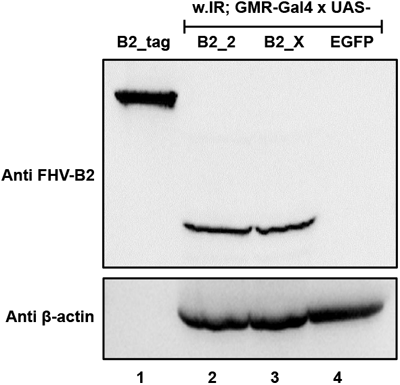
Western blot confirming expression of FHV-B2 protein. Lane 1, MBP-tagged B2 protein expressed from bacteria (positive control). Lanes 2 and 3, Expression of FHV-B2 in GMR-*white*IR; GMR-GAL4; UAS-FHV (chromosome 2; VK1 or X chromosome; attP18) flies. Lane 4, EGFP-expressed from GMR-*white*IR; GMR-GAL4; UAS-GFP (VK-1) flies (negative control). Anti-β-Actin was used to show equal loading.

## REFERENCES

[1] S. Dong, G. Dimopoulos, Aedes aegypti Argonaute 2 controls arbovirus infection and host mortality., Nature Communications 14 (2023) 5773. 10.1038/s41467-023-41370-y.

[2] Z. Guo, Y. Li, S.-W. Ding, Small RNA-based antimicrobial immunity, Nature Reviews Immunology 19 (2019) 31–44. 10.1038/s41577-018-0071-x.

[3] S.H. Merkling, A.B. Crist, A. Henrion-Lacritick, L. Frangeul, E. Couderc, V. Gausson, H. Blanc, A. Bergman, A. Baidaliuk, O. Romoli, M.-C. Saleh, L. Lambrechts, Multifaceted contributions of Dicer2 to arbovirus transmission by Aedes aegypti., Cell Reports 42 (2023) 112977. 10.1016/j.celrep.2023.112977.

[4] G.H. Samuel, T. Pohlenz, Y. Dong, N. Coskun, Z.N. Adelman, G. Dimopoulos, K.M. Myles, RNA interference is essential to modulating the pathogenesis of mosquito-borne viruses in the yellow fever mosquito Aedes aegypti, Proceedings of the National Academy of Sciences 120 (2023). 10.1073/pnas.2213701120.

[5] G.H. Samuel, Z.N. Adelman, K.M. Myles, Antiviral Immunity and Virus-Mediated Antagonism in Disease Vector Mosquitoes, Trends in Microbiology 26 (2018) 447–461. 10.1016/j.tim.2017.12.005.

[6] S.-W. Ding, Transgene Silencing, RNA Interference, and the Antiviral Defense Mechanism Directed by Small Interfering RNAs, Phytopathology® 113 (2023) 616–625. 10.1094/PHYTO-10-22-0358-IA.

[7] B.C. Bonning, M.-C. Saleh, The Interplay Between Viruses and RNAi Pathways in Insects, Annual Review of Entomology 66 (2021) 61–79. 10.1146/annurev-ento-033020-090410.

[8] W.-X. Li, S.-W. Ding, Mammalian viral suppressors of RNA interference, Trends in Biochemical Sciences 47 (2022) 978–988. 10.1016/j.tibs.2022.05.001.

[9] S.-W. Ding, O. Voinnet, Antiviral Immunity Directed by Small RNAs, Cell 130 (2007) 413–426. 10.1016/j.cell.2007.07.039.

[10] L.K. Johansen, J.C. Carrington, Silencing on the Spot. Induction and Suppression of RNA Silencing in the Agrobacterium -Mediated Transient Expression System, Plant Physiology 126 (2001) 930–938. 10.1104/pp.126.3.930.

[11] Y.S. Lee, R.W. Carthew, Making a better RNAi vector for Drosophila: use of intron spacers, Methods 30 (2003) 322–329. 10.1016/S1046-2023(03)00051-3.

[12] F. Li, S.-W. Ding, Virus Counterdefense: Diverse Strategies for Evading the RNA-Silencing Immunity, Annual Review of Microbiology 60 (2006) 503–531. 10.1146/annurev.micro.60.080805.142205.

[13] K.W.R. van Cleef, J.T. van Mierlo, M. van den Beek, R.P. van Rij, Identification of Viral Suppressors of RNAi by a Reporter Assay in Drosophila S2 Cell Culture, in: 2011: pp. 201–213. 10.1007/978-1-61779-037-9_12.

[14] B. Alberts, A. Johnson, L. J, et al, Molecular Biology of the Cell, 4th ed., Garland Science, 2002.

[15] K.S. Weiler, B.T. Wakimoto, Heterochromatin and gene expression in Drosophila., Annual Review of Genetics 29 (1995) 577–605. 10.1146/annurev.ge.29.120195.003045.

[16] H. Li, W.X. Li, S.W. Ding, Induction and Suppression of RNA Silencing by an Animal Virus, Science 296 (2002) 1319–1321. 10.1126/science.1070948.

[17] A. Nayak, B. Berry, M. Tassetto, M. Kunitomi, A. Acevedo, C. Deng, A. Krutchinsky, J. Gross, C. Antoniewski, R. Andino, Cricket paralysis virus antagonizes Argonaute 2 to modulate antiviral defense in Drosophila, Nature Structural & Molecular Biology 17 (2010) 547–554. 10.1038/nsmb.1810.

[18] R.P. van Rij, M.-C. Saleh, B. Berry, C. Foo, A. Houk, C. Antoniewski, R. Andino, The RNA silencing endonuclease Argonaute 2 mediates specific antiviral immunity in Drosophila melanogaster, Genes & Development 20 (2006) 2985–2995. 10.1101/gad.1482006.

[19] L.A. Perkins, L. Holderbaum, R. Tao, Y. Hu, R. Sopko, K. McCall, D. Yang-Zhou, I. Flockhart, R. Binari, H.-S. Shim, A. Miller, A. Housden, M. Foos, S. Randkelv, C. Kelley, P. Namgyal, C. Villalta, L.-P. Liu, X. Jiang, Q. Huan-Huan, X. Wang, A. Fujiyama, A. Toyoda, K. Ayers, A. Blum, B. Czech, R. Neumuller, D. Yan, A. Cavallaro, K. Hibbard, D. Hall, L. Cooley, G.J. Hannon, R. Lehmann, A. Parks, S.E. Mohr, R. Ueda, S. Kondo, J.-Q. Ni, N. Perrimon, The Transgenic RNAi Project at Harvard Medical School: Resources and Validation, Genetics 201 (2015) 843–852. 10.1534/genetics.115.180208.

[20] B.D. Pfeiffer, T.-T.B. Ngo, K.L. Hibbard, C. Murphy, A. Jenett, J.W. Truman, G.M. Rubin, Refinement of Tools for Targeted Gene Expression in Drosophila, Genetics 186 (2010) 735–755. 10.1534/genetics.110.119917.

[21] K.J.T. Venken, Y. He, R.A. Hoskins, H.J. Bellen, P[acman]: A BAC Transgenic Platform for Targeted Insertion of Large DNA Fragments in D. melanogaster, Science 314 (2006) 1747– 1751. 10.1126/science.1134426.

[22] M. Gaszner, G. Felsenfeld, Insulators: exploiting transcriptional and epigenetic mechanisms, Nature Reviews Genetics 7 (2006) 703–713. 10.1038/nrg1925.

[23] R.R. Roseman, V. Pirrotta, P.K. Geyer, The su(Hw) protein insulates expression of the Drosophila melanogaster white gene from chromosomal position-effects., The EMBO Journal 12 (1993) 435–442. 10.1002/j.1460-2075.1993.tb05675.x.

[24] M. Markstein, C. Pitsouli, C. Villalta, S.E. Celniker, N. Perrimon, Exploiting position effects and the gypsy retrovirus insulator to engineer precisely expressed transgenes, Nature Genetics 40 (2008) 476–483. 10.1038/ng.101.

[25] A.H. Brand, N. Perrimon, Targeted gene expression as a means of altering cell fates and generating dominant phenotypes, Development 118 (1993) 401–415. 10.1242/dev.118.2.401.

[26] M. Freeman, Reiterative Use of the EGF Receptor Triggers Differentiation of All Cell Types in the Drosophila Eye, Cell 87 (1996) 651–660. 10.1016/S0092-8674(00)81385-9.

[27] B.A. Hay, T. Wolff, G.M. Rubin, Expression of baculovirus P35 prevents cell death in Drosophila., Development (Cambridge, England) 120 (1994) 2121–9. 10.1242/dev.120.8.2121.

[28] S.M. Mackenzie, M.R. Brooker, T.R. Gill, G.B. Cox, A.J. Howells, G.D. Ewart, Mutations in the white gene of Drosophila melanogaster affecting ABC transporters that determine eye colouration, Biochimica et Biophysica Acta (BBA) - Biomembranes 1419 (1999) 173–185. 10.1016/S0005-2736(99)00064-4.

[29] R. Aliyari, Q. Wu, H.-W. Li, X.-H. Wang, F. Li, L.D. Green, C.S. Han, W.-X. Li, S.-W. Ding, Mechanism of induction and suppression of antiviral immunity directed by virus-derived small RNAs in Drosophila., Cell Host & Microbe 4 (2008) 387–97. 10.1016/j.chom.2008.09.001.

[30] J.A. Chao, J.H. Lee, B.R. Chapados, E.W. Debler, A. Schneemann, J.R. Williamson, Dual modes of RNA-silencing suppression by Flock House virus protein B2, Nature Structural & Molecular Biology 12 (2005) 952–957. 10.1038/nsmb1005.

[31] A. Lingel, B. Simon, E. Izaurralde, M. Sattler, The structure of the flock house virus B2 protein, a viral suppressor of RNA interference, shows a novel mode of double-stranded RNA recognition, EMBO Reports 6 (2005) 1149–1155. 10.1038/sj.embor.7400583.

[32] X.-H. Wang, R. Aliyari, W.-X. Li, H.-W. Li, K. Kim, R. Carthew, P. Atkinson, S.-W. Ding, RNA Interference Directs Innate Immunity Against Viruses in Adult Drosophila, Science 312 (2006) 452–454. 10.1126/science.1125694.

[33] J. Bischof, R.K. Maeda, M. Hediger, F. Karch, K. Basler, An optimized transgenesis system for Drosophila using germ-line-specific ΦC31 integrases, Proceedings of the National Academy of Sciences 104 (2007) 3312–3317. 10.1073/pnas.0611511104.

[34] B. Berry, S. Deddouche, D. Kirschner, J.-L. Imler, C. Antoniewski, Viral suppressors of RNA silencing hinder exogenous and endogenous small RNA pathways in Drosophila., PloS One 4 (2009) e5866. 10.1371/journal.pone.0005866.

[35] D.C. Baulcombe, The Role of Viruses in Identifying and Analyzing RNA Silencing., Annual Review of Virology 9 (2022) 353–373. 10.1146/annurev-virology-091919-064218.

[36] S. Liu, Y. Han, W.-X. Li, S.-W. Ding, Infection Defects of RNA and DNA Viruses Induced by Antiviral RNA Interference, Microbiology and Molecular Biology Reviews 87 (2023). 10.1128/mmbr.00035-22.

[37] H.M. Thorpe, M.C.M. Smith, In vitro site-specific integration of bacteriophage DNA catalyzed by a recombinase of the resolvase/invertase family, Proceedings of the National Academy of Sciences 95 (1998) 5505–5510. 10.1073/pnas.95.10.5505.

[38] A.K. Gupta, S. Tatineni, Wheat streak mosaic virus P1 Binds to dsRNAs without Size and Sequence Specificity and a GW Motif Is Crucial for Suppression of RNA Silencing, Viruses 11 (2019) 472. 10.3390/v11050472.

